# The Effect of Musical Groove on Sensorimotor Network Cortical Dynamics during Active Tapping and Passive Listening

**DOI:** 10.1101/2025.11.07.687225

**Authors:** Tommaso Zampini, Jessica M Ross, Petr Janata, Ramesh Balasubramaniam

## Abstract

Musical groove, defined as the urge to move in response to rhythmic stimuli, drives spontaneous entrainment and engagement with music. Recent studies have investigated the activation of the sensorimotor network (SMN), focusing on the µ rhythm, an oscillatory activity in the alpha (8-12 Hz) and beta (16-24 Hz) ranges that reflects motor system engagement. While µ modulation differs between passive listening and active movement, it remains unclear whether these responses are shaped by the level of musical groove. Clarifying this relationship is crucial for understanding how neural oscillations transform rhythmic perception into embodied movement and how auditory and motor systems interact to generate the spontaneous urge to move. The present study investigates SMN dynamics by examining µ modulation across two factors: mode of engagement - active motor entrainment (finger tapping) versus passive listening, and groove level - low versus high. Electroencephalography (EEG) recordings were used to assess SMN activity, with a focus on µ rhythm modulation in the alpha and beta frequency bands. Channel and cluster-based EEG analyses revealed stronger µ suppression during active tapping compared to passive listening, as well as an effect of musical groove. Self-reported groove ratings were not affected by the tapping task. Together, these findings highlight the modulation of the µ rhythm in the SMN, describing distinctive patterns of neural engagement during the experience of musical groove.

## Introduction

Groove refers to the compelling urge to move in synchrony with rhythmic music, a pleasurable phenomenon in which emotional engagement and bodily entrainment converge into a unified experience (Berliner, 1994; Janata et al., 2012; Pressing, 2002; Madison, 2006). It is often associated with a pleasurable state where the engagement with music feels effortless, fostering a sense of connection to the music itself and sometimes to other listeners or performers (Janata et al., 2012; Witek et al., 2014). Groove is widely considered an example of musical embodiment, where cognitive and affective processes manifest in the form of coordinated, temporally aligned movement (Etani et al., 2024).

The embodied experience of groove can emerge not only during active movement to music, such as in sensorimotor synchronization (SMS) tasks (Repp & Su, 2013), but also during passive listening, indicating that the neural systems supporting groove may be engaged independently of overt motor behavior. Using functional magnetic resonance imaging (fMRI), Chen et al. (2008) demonstrated that rhythmic processing activates brain regions within the sensorimotor network (SMN) that are also implicated in movement executions, even in the absence of movement. This SMN includes the supplementary motor area, premotor cortex, cerebellum, and basal ganglia. These results have been replicated numerous times (Gordon et al., 2018) and suggest that audio-motor coupling is a feature of both active and passive engagement with rhythm. Because groove fundamentally involves the integration of auditory and motor processes, these findings imply that its underlying mechanisms may operate during rhythm perception alone. Thus, to fully understand groove, a deeper investigation of the neural dynamics that operate across both active and passive modes of engagement is required.

Recent EEG studies have shown that passive music listening elicits distinct patterns of oscillatory activity across the alpha, beta, and gamma bands (Iversen et al., 2009; Fujioka et al., 2012; Ross et al., 2013; Comstock et al., 2021; Rosso et al., 2025). These oscillatory changes in the alpha activity are measured over the SMN, and include a peculiar signature called the “mu (µ) rhythm” which reflects motor system engagement (Pfurtscheller & Da Silva, 1999). The µ rhythm is generally represented by an amplitude peak in the alpha band (8 to 12 Hz) and first harmonic in the beta band (16 to 24 Hz). Additionally, Pfurtscheller & Neuper (1994) have shown that the µ rhythm is modulated by the mode of engagement with music: during voluntary movement tasks, µ power is suppressed over task-relevant motor areas but is increased over motor regions uninvolved in the action.

In contrast to methods that rely on SMS (active tapping tasks), recording EEG during passive listening allows us to explore neural dynamics in motor regions that are not directly related to executed action. Ross et al. (2022) showed music-related µ modulation in the absence of overt movements, supporting motor inhibition during listening. This study recorded EEG during silence without movement, with overt movements (foot/hand), and during music listening without movement. When recorded over the hand area of the primary motor cortex, the hand movement condition showed µ power suppression in the alpha band, while foot movement and passive music listening conditions showed enhanced µ power. Ross et al (2022) interpreted this increased µ power during music listening to reflect suppression of an urge to move in response to music.

Here, we investigated this phenomenon further by looking at the role of groove, and whether the degree of µ modulation changes across different levels of perceived groove. While neural oscillations during music listening have been studied, groove-specific neural responses during passive listening remain limited. Most studies of groove have focused on active SMS tasks, leaving the neural mechanisms of groove perception during passive listening largely unexplored. The present study addresses this gap by examining µ rhythm activity in the SMN during passive listening to music with varying groove levels, and by comparing these responses to active tapping conditions. We hypothesized that µ activity, representing movement suppression during passive listening to music, would be sensitive to groove level, possibly showing increased µ power for high groove music. Additionally, we explored whether subjective groove ratings corresponded to these neural activations, to develop insight into the relationship between perceived groove and SMN activation.

## Methods

### Participants

All participants (*n* = 23; mean age = 20.7, ± 2.48 years) were UC Merced undergraduate students who reported normal hearing and had no history of neurological, auditory, or motor impairments. This study was conducted in accordance with the Declaration of Helsinki and was reviewed and approved by the UC Merced Institutional Review Board (IRB). Prior to participation, each participant provided written informed consent.

An *a priori* power analysis was conducted using G*Power (Version 3.1.9.6), (Faul et al., 2007), based on effect sizes derived from Ross et al. (2022) who reported significant task-related modulation of µ rhythm activity across different Independent Component Analysis (ICA) source clusters. Using observed *F-statistics* reported in the author’s analysis, we first converted these into partial eta-squared (*η²ₚ)* and subsequently, into Cohen’s *f.* The left hemisphere source cluster alpha band spectral analysis reported *F*(3,140) = 8.63 (*η²ₚ* = 0.15, *f*= 0.43), left beta band *F*(3,212) = 23.22 (*η²ₚ* = 0.25, *f*= 0.57), midline alpha *F*(3,140) = 9.57 (*η²ₚ* = 0.17, *f* = 0.41), and right cluster alpha *F*(3,140) = 8.01 (*η²ₚ* = 0.15, *f* = 0.45). Based on these values, *a priori* sample size estimated for a repeated-measures ANOVA (two conditions and two levels, α = 0.05, power = 0.95) ranged from 14 (left beta) to 22 (midline alpha). We do not have effect sizes to estimate required sample size to detect the main effect of groove level, so we chose the sample size to minimally replicate the main task-related effects reported by Ross et al. (2022) regardless of brain region. We determined that we should have a minimum of *n*=22. We recruited a sample of *n* = 23 participants, which for the smallest effect size reported in Ross et al (*f* = 0.41) would achieve a power of 0.96 and for the left alpha condition (*f* = 0.43) would achieve a power of 0.98.

### Experimental Design

The experimental stimuli were controlled using Paradigm software™ (version 2.5.0.68). We included two main experimental conditions (passive listening and active tapping), along with a baseline rest condition. These conditions assessed both subjective and neural responses to groove perception under conditions of covert and overt motor engagement. The tapping and no-tapping conditions were presented in separate blocks. The baseline condition was repeated five times (twice at the beginning of the session, once between the no-tapping and tapping blocks, and twice again at the end) to ensure a stable neural baseline. During the no-tapping condition, participants listened passively to randomized music excerpts without performing any movement. An experimenter monitored participants throughout the session to ensure they remained still and did not engage in any overt movements, in line with previous observations that minor spontaneous motion can occur even during passive listening (Janata et al., 2012). This condition allowed for the measurement of neural entrainment in the absence of overt motor activity. Finally, in the tapping condition, participants were instructed to actively synchronize their finger taps to the beat of the music using their dominant index finger on a Makey Makey sensor pad (Collective & Shaw, 2012). This task was intended to engage the sensorimotor system through active listening. Across the total set of trials, each music excerpt was repeated five times, in each of the no-tapping and tapping conditions, to ensure consistency and reliability. In the baseline condition, participants fixated on a cross displayed at the center of the screen for 30 seconds while EEG data were recorded.

### Musical stimuli

Six music excerpts were selected for the experiment, each lasting 30 seconds (Table 1): three high-groove (HG) and three low-groove (LG) excerpts. These were chosen based on groove ratings reported by Janata et al. (2012). Excerpts were matched for vocals, meter, and beats per minute (BPM) to control for some possible effects of musical features on µ power (Stupacher et al., 2013).

**Table 1.**
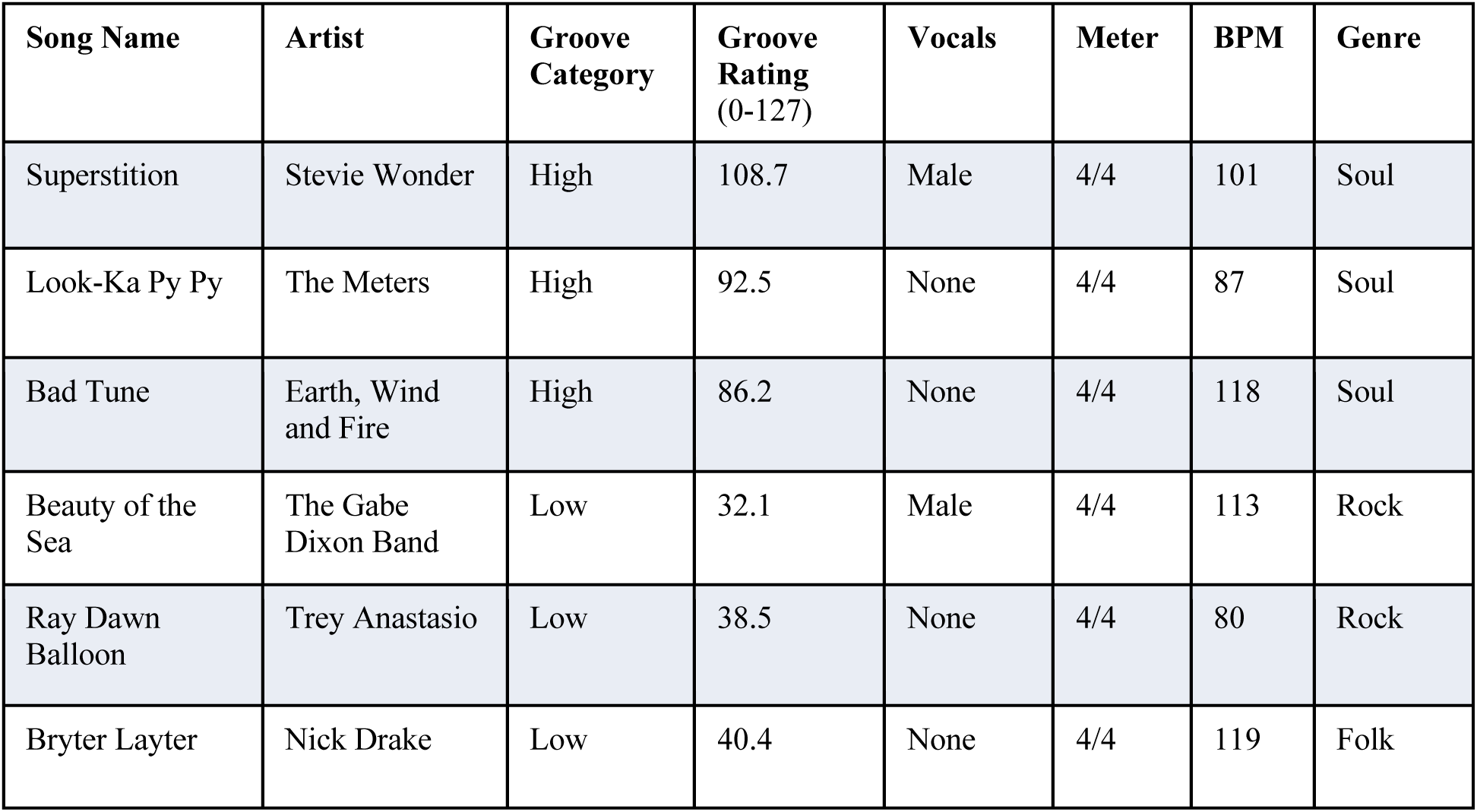
The 6 musical clips used as stimuli, information about their musical features and genre, and groove ratings (Janata et al., 2012). High- and low-groove clips were matched for similar vocals, meter, and tempo (BPM; Stupacher et al., 2013).

### Behavioral Tasks

We collected self-reported groove ratings through Paradigm software and analyzed them in R Studio. After each excerpt, in both passive and active conditions, participants verbally rated their sense of groove for the just-heard piece of music on a 5-point Likert scale displayed on the screen. They provided one of the following ratings, either by number or by words, to the experimenter in response to “*Please report your perceived groove verbally in response to the previous excerpt.*”: *1= Not at all, 2= A little, 3= Moderately, 4= A lot, 5 =Extremely*. In the active tapping condition, participants were additionally instructed to synchronize their finger taps to the beat of the music (as best as they could) using their dominant index finger on a Makey Makey sensor pad (Collective & Shaw, 2012). Tapping data were collected and tap times were measured respectively for both LG and HG music.

### Electroencephalography

#### EEG Data Collection

Before starting the experiment, participants were briefly familiarized through verbal instructions with the tapping task and EEG setup. EEG data were recorded using the ANT Neuro Waveguard 32-channel EEG system positioned using the international 10-20 system and with sintered Ag/AgCl electrodes (Asalab amplifier and Asalab software). The EEG cap was positioned relative to fiducials and with channel Cz at the midpoint between the nasion and inion. Data were sampled at 1024 Hz. If any electrodes showed high impedance before data collection (±5 kOhm), the participant’s hair was parted, and the scalp was gently rubbed in those locations to improve signal quality. Trial onsets were marked in the EEG recording by event triggers sent from Paradigm to Asalab.

#### EEG Data Cleaning

EEG data were cleaned and analyzed using MATLAB R2024b (MathWorks, Natick, MA) and the EEGLAB toolbox (Delorme & Makeig, 2004). Electrode locations were first loaded (*pop_chanedit*) and EEG data were then downsampled to 256 Hz before filtering the data (*pop_eegfilt*) using a 2 Hz high-pass and 50 Hz low-pass filter (Winkler et al., 2015; Ross et al., 2022). Noisy or drifting channels were manually identified and interpolated (*pop_interp;* mean ± standard deviation number of channels removed = 3.21±1.63) and data were subsequently re-referenced (*pop_reref*) to a sample-by-sample all-channels average. Continuous EEG data were segmented into multiple epochs of 32 seconds each corresponding to 2 seconds before and 30 seconds after the music excerpt’s onsets (extracted using event codes for the analysis). Epochs containing aberrant activity were excluded according to conservative criteria (Delorme et al., 2007; Ross et al., 2022), with an amplitude threshold set at ±500 µV (*pop_eegtresh*) and a probability test (*pop_jointprob*) for single (±6 SD threshold) and for multi-channel (±2 SD threshold) activities (Delorme et al., 2007). ICA (*runica*) was performed (*eeglab_ica*) on each participant’s preprocessed data (Bell & Sejnowski, 1995). ICA decomposes multichannel EEG into maximally independent components (ICs), allowing scalp-recorded signals to be expressed as a sum of underlying brain sources and non-neural artifacts (Burunat et al., 2017). Each IC can then be projected back to the scalp by combining its activation time course with its associated scalp map, which reflects its topographic pattern across the electrode montage (Makeig et al., 1997, 2004).

Each IC was visualized using the Viewprops plugin (*pop_viewprops*) and rejected if both the artifact classification exceeded 80% for eye, heart activities, channel noise, or muscle, and upon expert review the topoplots were consistent with this determination (mean ± standard deviation number of ICs removed = 4.8 ±2.6), (Delorme & Makeig, 2004).

Dipole fitting was conducted with alignment of channel locations to a head model based on the standard Montreal Neurological Institute (MNI) magnetic resonance (MR) image (*pop_dipfit_settings*). Single equivalent dipole fitting was then performed (*dipfit5.5)* and ICs with dipoles outside the head were removed using a residual variance <15% (*Fieldtrip* extension).

We used three channels over SMN for a channel level analysis (channels C3, Cz, C4) and also did an analysis in source space using a dipole clustering technique. For this clustering, we used *k*-means clustering, excluding outlier clusters (with dipole positions ≥3 SD from any of the cluster centroids), (Onton & Makeig, 2006; Ross et al., 2022). To maximize the number of subjects per cluster, we used three different cluster number optimization approaches (Calinski-Harabasz, Silhouette, and Davies-Bouldin) which suggested that between 12 and 14 clusters would be optimal. We therefore clustered into 14 clusters (plus 1 outlier cluster).

Of these 14 clusters, three µ-related clusters were identified by visual inspection that met a set of specific and pre-defined criteria following Pfurtscheller & McFarland (2012) and Ross et al. (2022). The criteria were that: 1) sources projected bilaterally to the scalp, exhibiting opposite polarity distributions across hemispheres and showed stronger projections over central compared to occipital regions; 2) the spectra had peaks in both the alpha and beta frequency bands, reflecting the harmonic structure of the µ rhythm; 3) all ICs in the cluster had similar dipole locations, covering left, central and right precentral gyri, specifically that the cluster centroid’s radial projection aligned with that region on the cortical surface. The clusters that were selected using these criteria contained no more than 2 ICs per participant and included ICs from 14-16 participants. Because our dipole localization relied on a standardized head model, these dipole locations are approximate to their actual cortical sources in each participant (Akalin & Makeig, 2013).

### Analyses

#### Groove Ratings

Raw groove ratings were imported from Excel files corresponding to the experimental conditions: HG tapping, HG no-tapping, LG tapping, and LG no-tapping. Ratings, originally collected on Paradigm as categorical labels (*Point1-Point5*), were converted to numeric values (1-5). Each trial was labeled by condition and aggregated into a single dataset. For descriptive purposes, mean ratings, standard deviations (*SD*), and standard errors (*SE*) were computed separately for each condition.

To test for task-related and groove-related main effects on the groove perception ratings, we performed a 2×2 repeated measures ANOVA with the factors task (tapping vs. no-tapping) and groove (HG vs. LG). Analyses were performed in R Studio using the *aov* function, and partial eta squared (*η²ₚ*) was reported as an effect size. Post-hoc pairwise comparisons were performed using estimated marginal means (*emmeans* package) with Bonferroni correction for multiple comparisons.

#### Tapping Measures

Tapping responses were collected via a Makey Makey sensor pad and recorded concurrently with the EEG signal. Tap timestamps were extracted for each participant and categorized as HG or LG depending on the musical excerpt presented during the corresponding trial. Beat onset times for each musical excerpt were computed using the *musicntwrk* Python package (Buongiorno Nardelli, 2019). Tap latencies were then aligned with the detected beat onsets to quantify synchronization performance. Latency and interval data were exported as Excel files and imported into R Studio where Inter-tap interval (ITI) variability, mean asynchrony and asynchrony variability (SD) were computed for each trial. To compare synchronization performance between HG and LG conditions, we conducted a series of Welch two-sample *t*-tests using the *t.test* function. For each measure, Cohen’s *d* was computed as an estimate of effect size using the *cohens_d* function (*effectsize* package).

#### EEG Measures

Spectral power analysis was conducted at both channel and cluster levels to examine µ signatures in the alpha (8-12 Hz) and beta (16-24 Hz) frequency bands (Pfurtscheller & Neuper, 1994; Wolpaw & Wolpaw, 2012). Mean absolute power spectral density (PSD) was calculated across epochs for the full 5-24 Hz range using the *spectopo* function (window size = 1024 samples, 0.25 Hz resolution, 50% overlap), and condition-specific spectra were plotted for visual comparison and figure generation. Spectral curves were smoothed with a 5-point moving average to reduce high-frequency fluctuations without obscuring relevant peaks.

Event-related spectral perturbations (ERSPs) were computed at both channel and cluster levels to capture time-frequency dynamics over sensorimotor regions. ERSPs were estimated using *newtimef* with a frequency range of 5–24 Hz, wavelet cycles = 3, and scaling factor = 0.8. Analyses used 200 frequency bins (*nfreqs*) and 200 time points (*ntimesout*) spanning a 32-s window (–2000 to 30,000 ms; ≈160 ms resolution per time point). A baseline segment of equal duration was used to normalize all conditions. At the cluster level, ERSPs were computed for each IC and then averaged across all ICs within the cluster to account for source-resolved neural dynamics.

Two experimental conditions and two levels were analyzed: active HG, active LG, passive HG, and passive LG. For statistical analyses, PSD values were extracted from the alpha (8-12 Hz) and beta (16-24 Hz) bands. Mean spectral power for task- and groove-related effects was compared using 2×2 repeated-measures ANOVAs with factors task (tapping vs. no-tapping) and groove (HG vs. LG). Analyses were performed in R Studio using the *aov* function, and partial eta squared (*η²ₚ*) was reported as an effect size. Post hoc pairwise comparisons were conducted using estimated marginal means (*emmeans* package) with Bonferroni correction for multiple comparisons.

## Results

### Groove Ratings

Participants’ self-reported groove ratings varied based on musical groove but not task (Fig. 1). The 2×2 repeated-measures ANOVA revealed a significant main effect of groove, *F*(1, 344) = 942.70, *p* < .001, *η²ₚ*= 0.73. Follow-up paired comparisons confirmed that HG stimuli were rated as more “groovy” than LG stimuli. There was no significant main effect of task, *F*(1, 344) = 0.37, *p* = .54, *η²ₚ*= 0.001, nor a significant task × groove interaction, *F*(1, 344) = 1.03, *p* = .31, *η²ₚ*= 0.003. Post hoc comparisons showed significantly higher groove ratings for HG compared to LG stimuli regardless of task, with no added groove enhancement from tapping.

**Figure 1.**
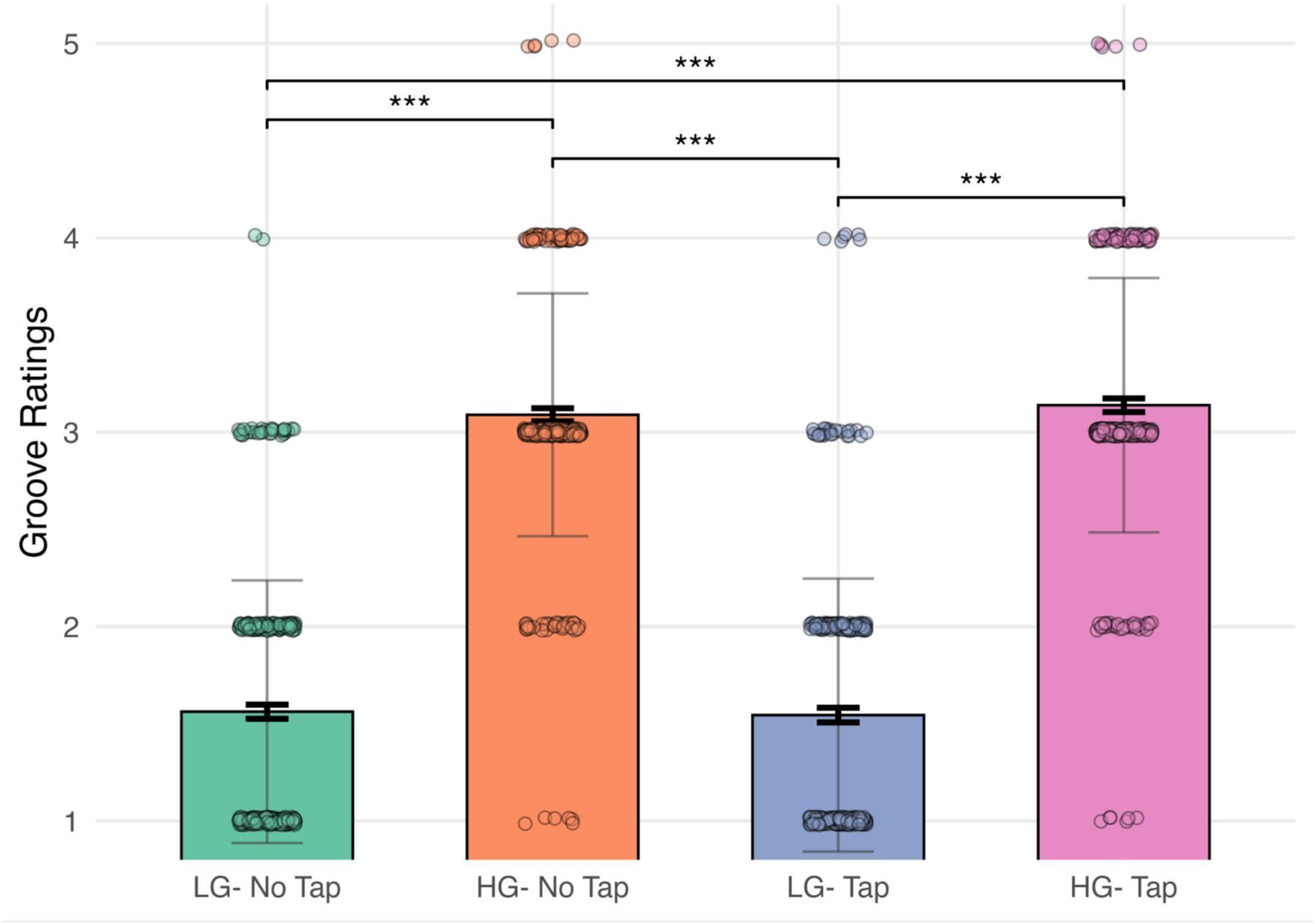
Self-reported groove ratings means, standard deviations and standard errors in tick lines (y axis - 1=Not at All, 2=A little, 3=Moderately, 4=A lot, 5=Extremely) for conditions (x axis - active HG, active LG, passive HG, passive LG). Brackets mark significant differences between condition pairs in the post hoc comparisons.

### Tapping

A series of Welch two-sample *t*-tests compared tapping performance between HG and LG music. Mean asynchrony (Fig. 2A) did not differ significantly between groove levels, *t*(673.50) = 1.85, *p* = .065, 95% *CI* [−0.0002, 0.0064], *d* = 0.14. However, inter-tap interval (ITI) coefficient of variation (Fig. 2B) was significantly lower during HG than LG music, *t*(671.55) = −2.94, *p* = .003, 95% *CI* [−0.12, −0.02], *d* = −0.23. Similarly, asynchrony variability (SD) (Fig. 2C) was significantly reduced for HG compared to LG music, *t*(668.71) = −3.15, *p* = .002, 95% *CI* [−0.0068, −0.0016], *d* = −0.24.

**Figure 2.**
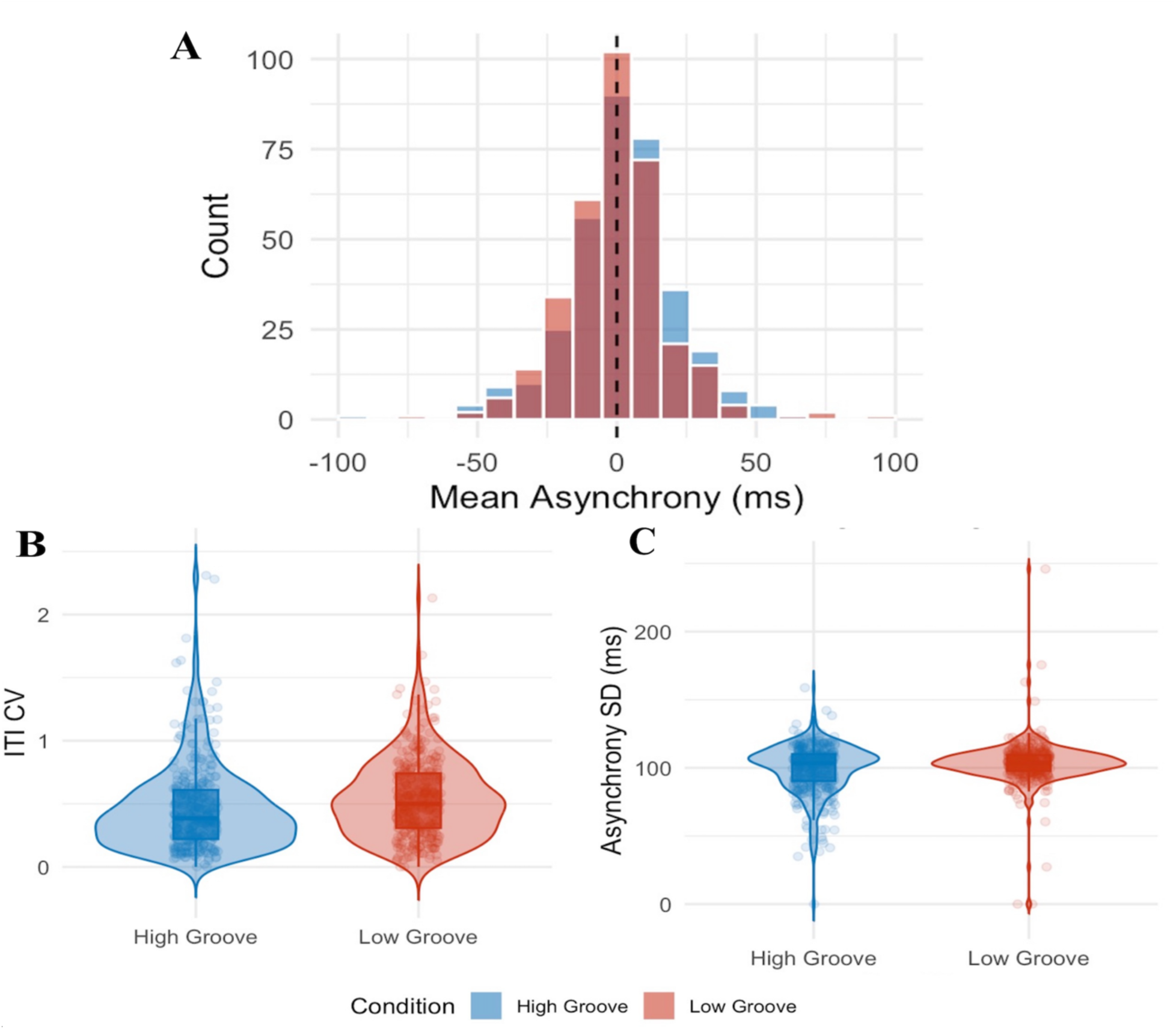
Beat-tap synchronization performance across groove conditions. **A**: Distribution of mean asynchrony - timing accuracy for HG (blue) and LG (red) music; dashed line indicates perfect synchrony with beats (0 ms). **B**: Inter-tap interval (ITI) coefficient of variation of tapping consistency (y axis) between HG and LG (x axis). **C**: Asynchrony SD - timing precision (ms; y axis) between conditions (x axis).

### EEG Channels

The spectral plot of C3 revealed visible peaks in the alpha (∼12 Hz) and beta (∼18 Hz) bands, reflecting μ-rhythm related activity (Appendix Fig 1A). Qualitatively, time-frequency plots appeared to support this pattern, showing pronounced event-related desynchronization (ERD) during active conditions (Appendix Fig. 1B). In the spectra, both active HG and LG elicited greater μ suppression than passive listening, suggesting greater motor engagement during active listening. These patterns are consistent with μ suppression during movement reported in the literature. This suppression was most evident at C3, compared to Cz and C4. The spectral plot of Cz revealed similar alpha and beta band peaks (∼10 Hz and ∼17 Hz), though differences in power between conditions was less clear than at C3 (Appendix Fig. 2A). Time-frequency plots supported ERD in active conditions, but showed a reduced trend compared to C3 (Appendix Fig. 2B). For C4, both spectra (peaks at ∼ 10 Hz and ∼18 Hz), (Appendix Fig. 3A) and ERSP plots showed similar patterns of alpha and beta activity with μ suppression during active listening (Appendix Fig. 3B). Across all channels, active HG (blue) consistently showed a stronger alpha-band suppression than active LG (red), while passive LG (purple) tended to show higher alpha power than passive HG (yellow), particularly at C3. A 2×2 repeated-measures ANOVA confirmed these effects (Appendix Tab. 1; Appendix Fig.4).

### EEG Clusters

#### Left Cluster

The left IC cluster (18 ICs from 14 participants) centroid was at Talairach coordinates (-37, -19, 49), (MRIcon, v1.0.20190902). The spectral analysis revealed visible peaks in the alpha (∼11 Hz) and beta (∼17 Hz) bands (Fig. 3C). Both HG and LG conditions in the active task appeared to produce µ rhythm suppression compared to passive listening tasks and the baseline (Fig. 6), reflecting motor-related desynchronization during tapping. Within the active tasks, alpha and beta power appeared different between HG and LG, suggesting distinct motor responses. ERSP plots supported more µ desynchronization in active than passive conditions, particularly in the alpha band (Appendix Fig. 5A).

**Figure 3.**
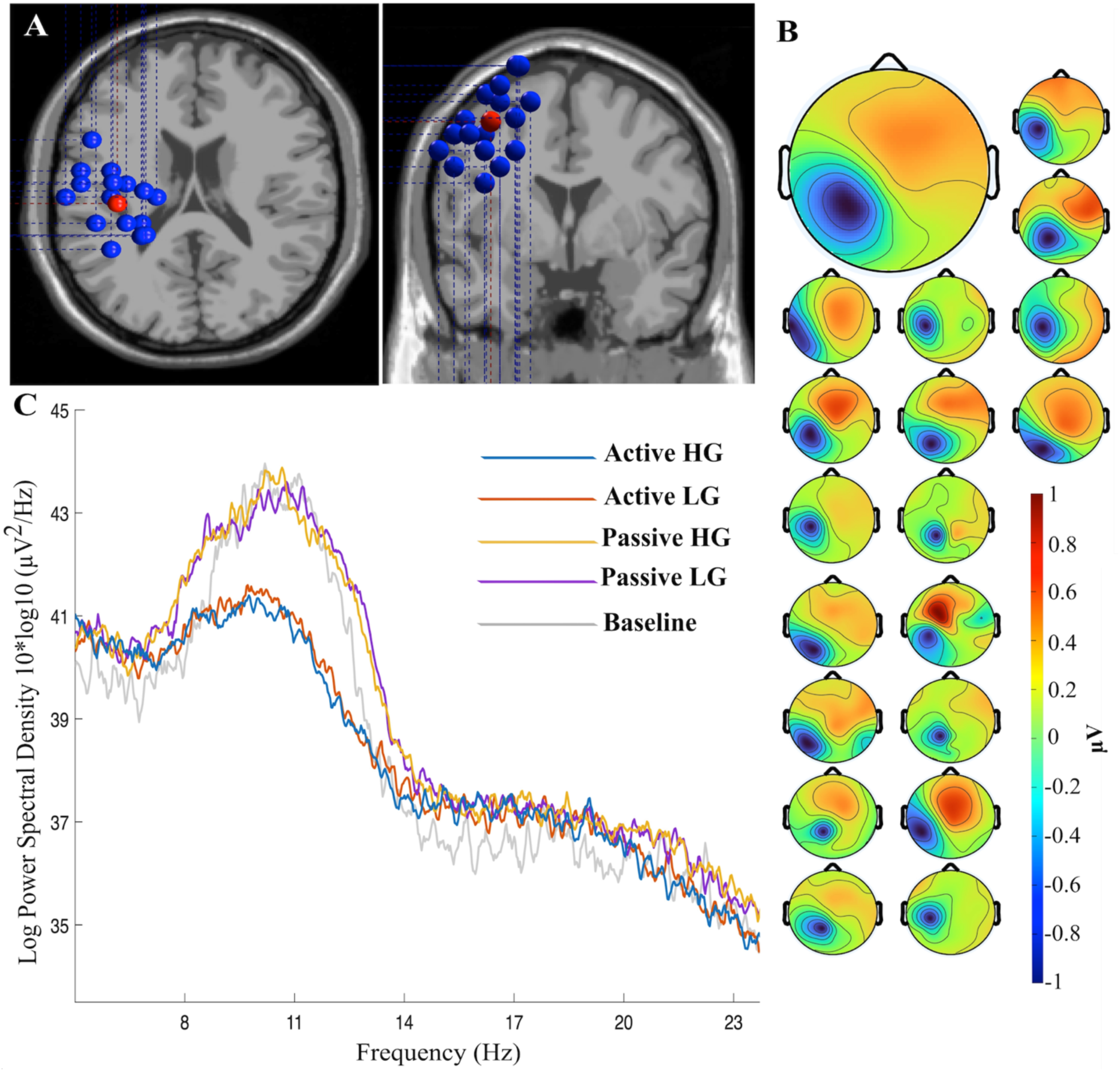
Left µ rhythm cluster (18 independent components from 14 participants). **A**: Individual dipoles (in blue) and centroid (in red) localized to Talairach (-37, -19, 49), approximately BA4. **B:** Average cluster scalp map and all independent components in the cluster. **C:** Mean log power spectra in the four experimental conditions + baseline. µ-related activity in alpha detected at ∼10-12 Hz; beta-band peaks (∼17 Hz) comprise µ harmonic and other beta-band activity.

Our 2×2 repeated-measures ANOVA supported that these patterns were significant. In the alpha band, a main effect of task was observed (*F*(1,32) = 427.64, *p* < .001, *η²p* = 0.93), along with a main effect of groove (F(1,32) = 122.60, p < .001, *η²ₚ* = 0.79) and a robust interaction between task and groove (*F*(1,32) = 46.92, *p* < .001, *η²ₚ*= 0.59). In the beta band, there were also main effects of task (*F*(1,64) = 72.86, *p* < .001, *η²ₚ*= 0.53), groove (*F*(1,64) = 75.11, *p* < .001, *η²ₚ*= 0.54), and an interaction between task and groove (*F*(1,64) = 13.12, *p* < .001, *η²ₚ*= 0.17).

#### Midline Cluster

The midline cluster (22 ICs from 17 participants) centroid was at Talairach coordinates (1, -14, 50), (MRIcon, v1.0.20190902). Spectral plots showed a slightly different pattern of results than the left hemisphere cluster (Fig. 4C). There appears to have been alpha suppression that was greater during active listening than during passive listening, though groove-specific patterns were less pronounced than in the left cluster. The plots show that in the beta band there appears to be an increase in power compared to the baseline for all of the conditions. Specifically, only passive listening conditions appear different from the baseline. ERSP plots support these patterns, showing a sustained power for passive listening tasks (Appendix Fig. 5B).

**Figure 4.**
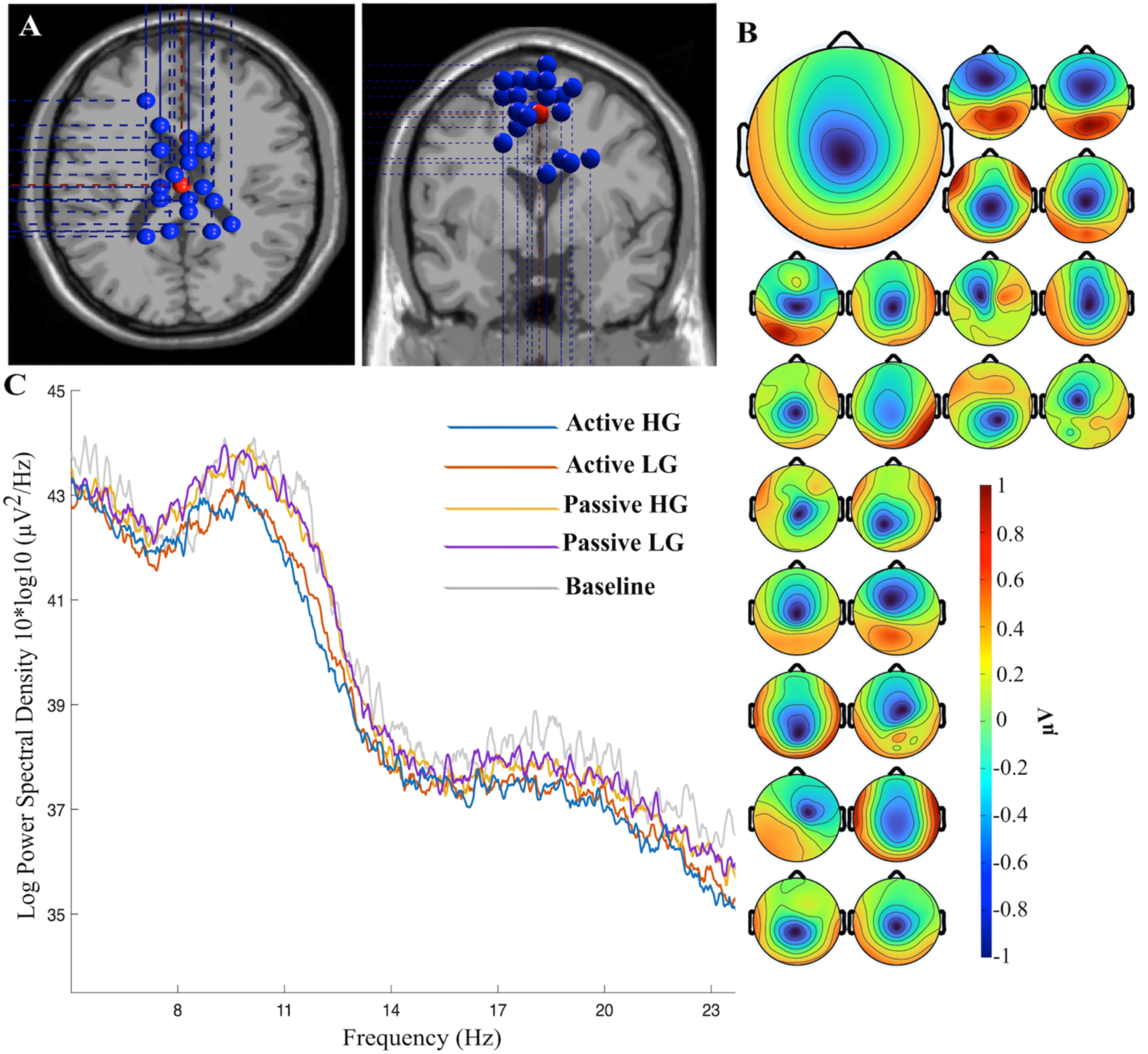
Midline µ rhythm cluster (22 independent components from 17 participants). **A**: Individual dipoles (in blue) and centroid (in red) localized to Talairach (1, -14, 50), approximately SMA. **B**: Average cluster scalp map and all independent components in the cluster. **C:** Mean log power spectra in the four experimental conditions + baseline. µ-related activity in alpha detected at ∼ 10 Hz; beta-band peaks (∼17 Hz) comprise µ harmonic and possibly other beta-band activity.

For the alpha band, the ANOVA revealed a main effect of task (*F*(1,32) = 482.30, *p* < .001, *η²p* = 0.94) and a main effect of groove (*F*(1,32) = 7.76, *p* < .01, *η²ₚ*= 0.20), but no significant interaction between task and groove (*F*(1,32) = 0.02, *p* = .88, *η²ₚ*= 0.001). For the beta band, there were main effects of both task (*F*(1,64) = 362.81, *p* < .001, *η²ₚ*= 0.85) and groove (*F*(1,64) = 19.34, *p* < .001, *η²ₚ*= 0.23), but no interaction (*F*(1,64) = 0.03, *p* = .87, *η²ₚ*= 0.001).

#### Right Cluster

The right IC cluster (21 ICs from 16 participants) centroid was at Talairach coordinates (45, -11, 49), (MRIcon, v1.0.20190902). The spectral plots showed a similar pattern of brain changes when compared to the left hemisphere cluster (Fig. 5C). There appeared to be alpha band suppression that was larger in active conditions, and differences between HG and LG within the active tasks. In the beta band, all of the conditions showed a power increase compared to the baseline. ERSPs showed a consistent power increase for passive HG and LG compared to active listening tasks for the entire task (Appendix Fig. 5C).

**Figure 5.**
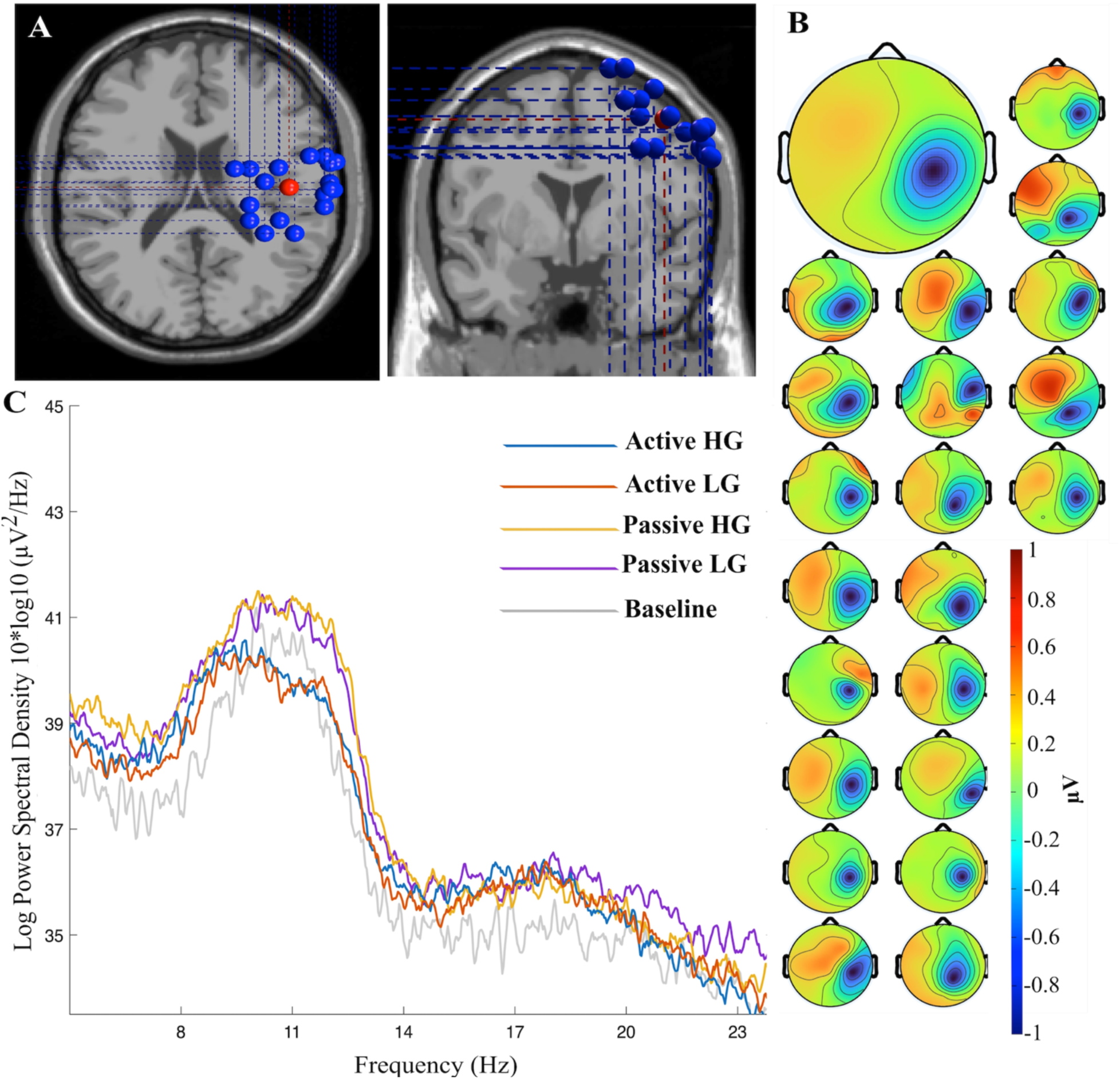
Right µ rhythm cluster (21 independent components from 16 participants). **A**: Individual dipoles (in blue) and centroid (in red) localized to Talairach (45, -11, 49), approximately BA4. **B:** Average cluster scalp map and all independent components in the cluster. **C:** Mean log power spectra in the four experimental conditions + baseline. µ-related activity in alpha detected at ∼10 Hz; beta-band peaks (∼18 Hz) comprise µ harmonic and possibly other beta-band activity.

**Figure 6.**
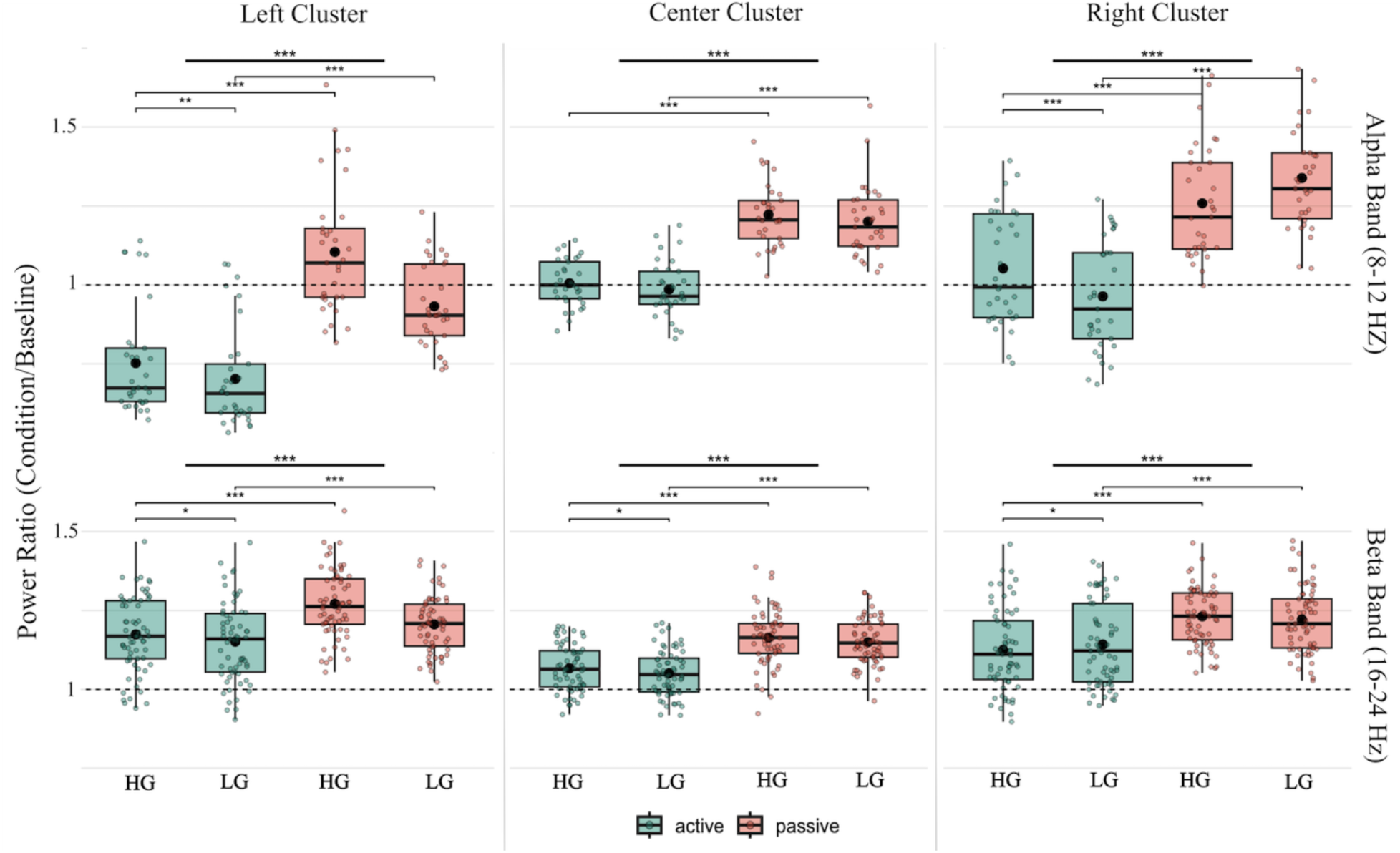
Ratios of mean power during active HG, active LG, passive HG and passive LG conditions to mean power in the baseline (ERD, this ratio < 1; ERS, this ratio > 1) for clusters (columns) at specific frequency bands (alpha - first row; beta - second row). Tick lines at the top mark significant differences between tasks (active in green vs. passive in pink); thin lines mark significant differences between pairs of conditions from the post hoc comparisons.

In the alpha band, the ANOVA revealed a significant main effect of task (*F*(1,32) = 406.68, *p* < .001, *η²ₚ*= 0.93), no main effect of groove (*F*(1,32) = 0.41, *p* = .52, *η²ₚ*= 0.01), but a strong interaction between task and groove (*F*(1,32) = 113.94, *p* < .001, *η²ₚ*= 0.78). In the beta band, a significant main effect was found for task (*F*(1,64) = 96.93, *p* < .001, *η²ₚ*= 0.60) but not for groove (*F*(1,64) = 0.65, *p* = .42, *η²p* = 0.01), and there was an interaction between task and groove (*F*(1,64) = 8.75, *p* < .01, *η²ₚ*= 0.12).

Taken together, these results show μ rhythm suppression during active listening in both the left and right clusters, with the left cluster showing the most robust task and groove-related modulation across both frequency bands. These analyses identified significant main effects and interactions, shedding light on how active synchronization versus passive listening interact with the degree of groove to modulate cortical oscillatory responses.

## Discussion

The present study provides insights into the role of groove and whether the degree of µ modulation in the SMN differs across levels of perceived groove. First, we examined whether our data replicate prior studies showing µ rhythm suppression with overt movement. Our results confirmed this, revealing significantly enhanced µ rhythm suppression during active tapping compared to passive listening, a finding that aligns with a body of research demonstrating the impact of motor execution on µ modulation (Pfurtscheller & Neuper, 1994; Pfurtscheller & Da Silva, 1999; Ross et al., 2022; Stegemöller et al., 2018, 2021). In addition to a suppression of µ during movement, finding the opposite pattern during passive listening highlights the dynamic role of motor engagement during listening. The increased µ power during passive listening suggests motor suppression, where the system is in a state of motor readiness that is primed for movement, but overt action is inhibited. This result may align with theoretical models such as Action Simulation for Auditory Prediction (ASAP; Patel & Iversen, 2014), which posits that the motor system internally simulates actions during rhythm perception. Therefore, these oscillatory patterns reflect sensorimotor processing associated with the task: suppression during active motor engagement during tapping and increased µ power during a listening state in which movement is inhibited. While these patterns could potentially relate to neural entrainment to the beat, further analyses (e.g., phase-locking) would be necessary to establish such a connection.

We also hypothesized that µ modulations would be sensitive to the music’s groove level and that subjective groove ratings would correspond to these neural signatures of SMN activity. Our data show that µ power was significantly modulated by whether the stimuli were HG or LG music, even in the absence of overt movement. Differences in µ power between HG and LG music within the same motor tasks offer evidence that groove directly influences this neural marker of entrainment. Our behavioral tapping results further supported this relationship. While mean asynchrony did not differ between HG and LG conditions, ITI and asynchrony variability showed significant enhancements for HG music. Specifically, HG music reduced ITI and asynchrony SD indicating more regular motor output and reduced beat-to-beat timing fluctuations. These results suggest that groove may strengthen internal timekeeping and stabilizing rhythmical motor outputs, rather than improving beat detection per se, consistent with theories proposing that groove primarily shapes motor planning and predictive timing rather than perceptual-motor coupling accuracy (Janata et al., 2012; Madison, 2006; Witek et al., 2014). These behavioral findings complement the observed µ rhythm modulations, suggesting that groove-related enhancements in motor stability may be mirrored at the neural level.

As for groove ratings, participants rated HG and LG excerpts significantly differently within the same motor task, yet no significant differences were found between passive and active listening conditions. Despite a lack of significant distinction between covert and overt motor entrainment, previously reported by Spiech et al., 2022, this evidence aligns with findings showing that groove can be influenced by musical features, such as moderate syncopation, prominent bass frequencies, and pronounced beat salience (Janata et al., 2012; Hove et al., 2014; Stupacher et al., 2022). Behavioral and neuroscientific evidence suggests that groove may also emerge when rhythmic patterns strike a balance between predictability and surprise (Witek et al., 2014; Vuust & Makeig, 2014; Stupacher et al., 2022), and that passive listening engages sensorimotor networks as well as reward-related regions, including the basal ganglia (Gold et al., 2019; Matthews et al., 2020). All of these elements enhance rhythmic stability while preserving a degree of temporal tension, a dynamic that appears to underlie the compulsion to move. Our findings may support that the tension between rhythmic predictability and flexibility is linked to movement planning. While these predictive mechanisms facilitate overt entrainment during SMS tasks (Repp & Su, 2013; Iversen & Balasubramaniam, 2016), the nature of covert motor system engagement is less well-understood (Ross & Balasubramaniam, 2022). For instance, a recent study by Cossavella et al. (2025) showed that enhanced µ power during passive listening correlated with poorer subsequent beat tapping performance, supporting an interpretation that alpha power (µ rhythm) might reflect inhibition of movement, which could decrease accuracy.

It should be noted that each dipole source cluster included a spread of dipole estimates. The observed variability among ICs within clusters likely reflects inherent limitations of the dipole localization method employed. Future investigations incorporating individual structural MRIs could substantially enhance the anatomical precision of dipole estimation and subsequent clustering. Such methodological refinements would improve the alignment between IC clusters and underlying cortical regions, thereby enabling a more accurate characterization of µ-specific modulations.

Additionally, it may be relevant to highlight the differential µ rhythm modulations observed across hemispheres. In particular, we found a more pronounced µ suppression in the left hemisphere than at the midline or in the right hemisphere. Since all participants in our sample tapped with the right hand, this left lateralization reflects contralateral organization of µ ERD/ERS. Although contralateral hand movement explains the left lateralization of µ changes, it is unknown whether handedness of the participants interacts with the size of the ERD/ERS. To further investigate this aspect, future studies should consider incorporating left-handed participants, to determine whether similar effects emerge in their contralateral hemisphere. Further, the interaction between hemispheres for regulation of excitatory and inhibitory influences on movement are not well-understood for µ ERD/ERS. Interhemispheric inhibition (IHI) contributes critically to fine motor control and tapping coordination, but the effects specifically on µ ERD/ERS should be directly studied, as well as the interaction with handedness.

It is unknown whether musicianship or other personal experience factors contribute to µ ERD/ERS and should be investigated in future work (Ridding et al., 2000; Kuo et al., 2019; Jakubowski et al., 2022; Spiech et al., 2022). Follow-up studies should examine how µ modulation relates to behavioral tapping data, cross-cultural differences in musical exposure, and individual differences in musical expertise (musicians vs. non-musicians; Dotov et al., 2021).

In conclusion, this work supports the view that groove is not merely a passive perceptual phenomenon, but an embodied state closely tied to the motor system, even during passive listening. Our results provide a critical link between the subjective experience of groove and the underlying neural signatures of the SMN, indicating that µ modulation may reflect embodied auditory prediction and motor readiness.

Given the strong coupling between groove perception and motor system engagement, these findings also suggest promising applications beyond music cognition, particularly in therapeutic contexts. Thus, the present work represents an initial but important step toward the study of groove-related neural dynamics for motor rehabilitation in clinical populations with movement impairments (Bella et al., 2015, 2018; Kang et al., 2023). Building on this foundation, future work could explore adaptive rhythmic stimulation, e.g. using an adaptive metronome (Fink et al., 2022), neurofeedback, or longitudinal intervention designs, opening the door to innovative music-based approaches in therapy and clinical rehabilitation.

## Data Availability Statement

### Author contributions

T.Z., and R.B. conceived and designed the research. T.Z. performed the experiment, analyzed the data, and prepared the figures. T.Z., and R.B. drafted the manuscript, which was then edited and revised by T.Z., J.M.R., P.J., and R.B. All authors interpreted the study’s results and approved the final manuscript version.

## Diversity in Citation Practices

Retrospective analysis of the citations in every article published in this journal from 2010 to 2021 reveals a persistent pattern of gender imbalance: Although the proportions of authorship teams (categorized by estimated gender identification of first author/last author) publishing in the Journal of Cognitive Neuroscience (JoCN) during this period were M(an)/M = .407, W(oman)/M = .32, M/ W = .115, and W/ W = .159, the comparable proportions for the articles that these authorship teams cited were M/M = .549, W/M = .257, M/ W = .109, and W/ W = .085 (Postle and Fulvio, JoCN, 34:1, pp. 1–3). Consequently, JoCN encourages all authors to consider gender balance explicitly when selecting which articles to cite and gives them the opportunity to report their article’s gender citation balance

## Appendix

**Appendix Table 1.** Results of 2×2 repeated-measures ANOVA on mean spectral power at ROI channels (C3, Cz, C4), showing main effects of Task (tapping vs. no-tapping), Groove (high vs. low groove), and their interaction in the alpha (8-12 Hz) and beta (16-24 Hz) bands. Reported values include F-statistics with degrees of freedom (*df*), p-values, and partial eta-squared (*η²ₚ).* Significance levels: *p* < .05 = *, *p* < .01 = **, *p* < .001 = ***, n.s. = not significant.

| Channel | Band | Effect | F(df) | p-value | $\eta^2_p$ | Significance |
| --- | --- | --- | --- | --- | --- | --- |
| C3 | Alpha | Task | F(1,32) = 133.62 | < .001 | 0.81 | *** |
| C3 | Alpha | Groove | F(1,32) = 5.39 | .027 | 0.14 | * |
| C3 | Alpha | Task × Groove | F(1,32) = 0.49 | .49 | 0.02 | n.s. |
| C3 | Beta | Task | F(1,64) = 112.65 | < .001 | 0.64 | *** |
| C3 | Beta | Groove | F(1,64) = 43.01 | < .001 | 0.42 | *** |
| C3 | Beta | Task × Groove | F(1,64) = 12.42 | < .001 | 0.16 | *** |
| Cz | Alpha | Task | F(1,32) = 8.72 | .006 | 0.21 | ** |
| Cz | Alpha | Groove | F(1,32) = 3.47 | .072 | 0.11 | n.s. |
| Cz | Alpha | Task × Groove | F(1,32) = 8.29 | .007 | 0.21 | ** |
| Cz | Beta | Task | F(1,64) = 362.64 | < .001 | 0.85 | *** |
| Cz | Beta | Groove | F(1,64) = 4.08 | .048 | 0.06 | * |
| Cz | Beta | Task × Groove | F(1,64) = 38.93 | < .001 | 0.38 | *** |
| C4 | Alpha | Task | F(1,32) = 175.84 | < .001 | 0.85 | *** |
| C4 | Alpha | Groove | F(1,32) = 0.29 | .60 | 0.01 | n.s. |
| C4 | Alpha | Task × Groove | F(1,64) = 1.49 | .23 | 0.04 | n.s. |
| C4 | Beta | Task | F(1,64) = 288.47 | < .001 | 0.82 | *** |
| C4 | Beta | Groove | F(1,64) = 36.63 | < .001 | 0.36 | *** |
| C4 | Beta | Task × Groove | F(1,64) = 8.52 | .005 | 0.12 | ** |

**Appendix Fig. 1.**
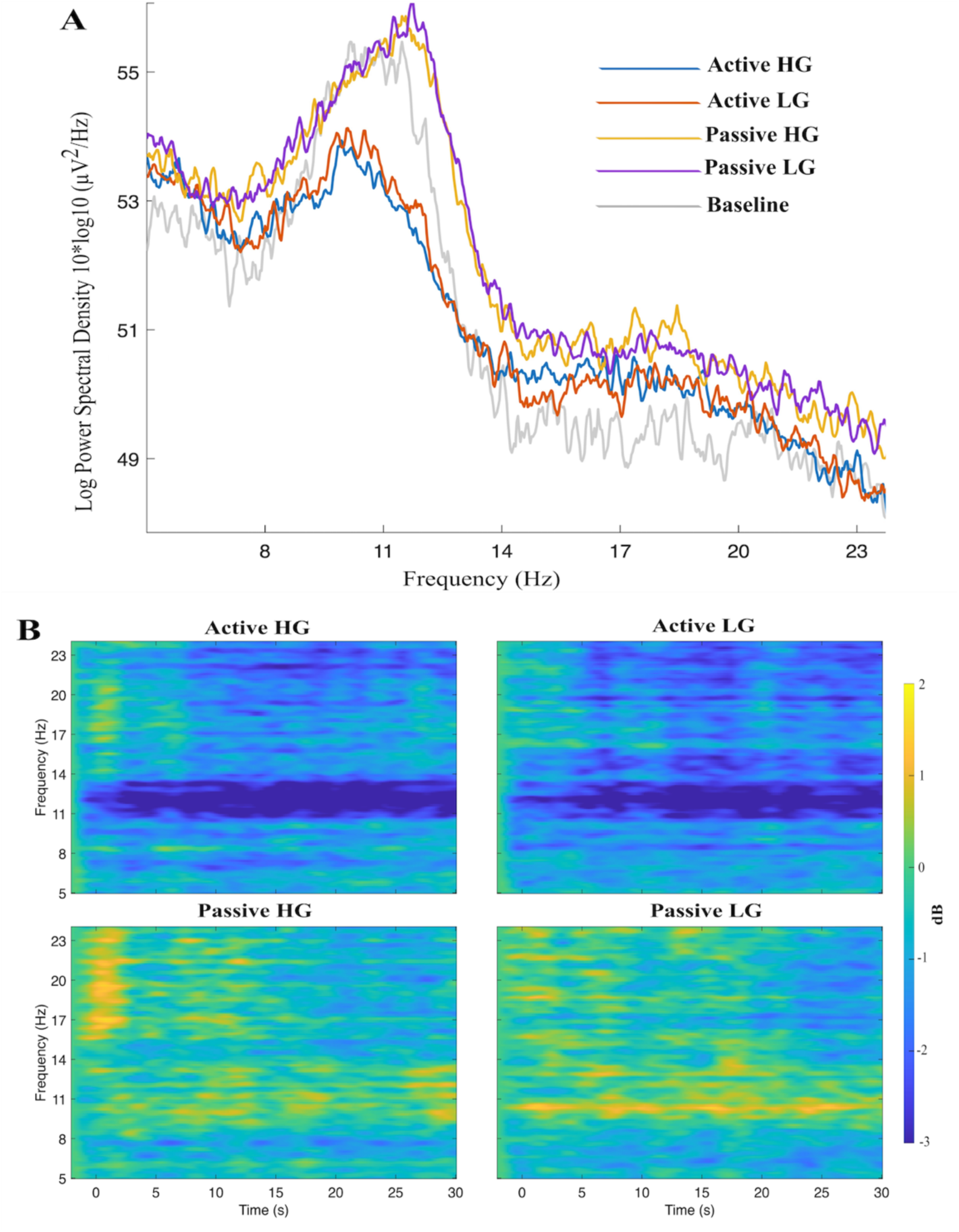
Spectral and ERSPs analyses of electrode C3. **A:** Mean log power spectra: power amplitude (reported in Log Power -y axis) at specific frequency bands (alpha and beta -x axis) for the four experimental conditions + baseline. µ-related activity in the alpha band was detected at ∼ 11-12 Hz; beta-band peaks (∼18 Hz) comprise µ harmonic and possibly other beta-band activity. **B**: Time-frequency plots: ERD (in blue) and ERS (in yellow) for alpha and beta (y axis) during the epoch duration (-2 to 30 sec from the stimulus onset -x axis).

**Appendix Fig. 2.**
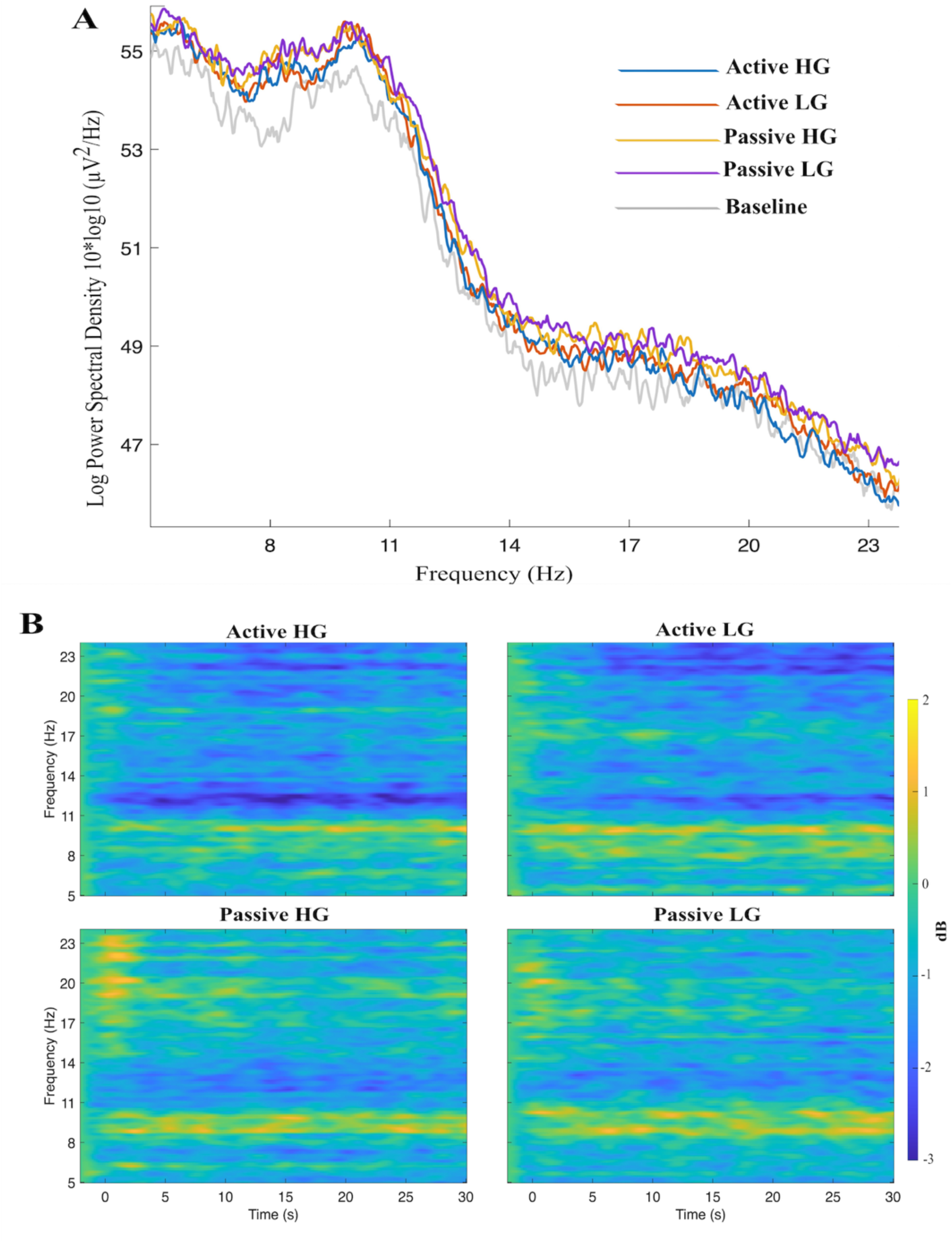
Spectral and ERSPs analyses of electrode Cz. **A:** Mean log power spectra: power amplitude (reported in Log Power -y axis) at specific frequency bands (alpha and beta -x axis) for the four experimental conditions + baseline. µ-related activity in the alpha band was detected at ∼10 Hz; beta-band peaks (∼17 Hz) comprise µ harmonic and possibly other beta-band activity. **B**: Time-frequency plots: ERD (in blue) and ERS (in yellow) for alpha and beta (y axis) during the epoch duration (-2 to 30 sec from the stimulus onset -x axis).

**Appendix Fig. 3.**
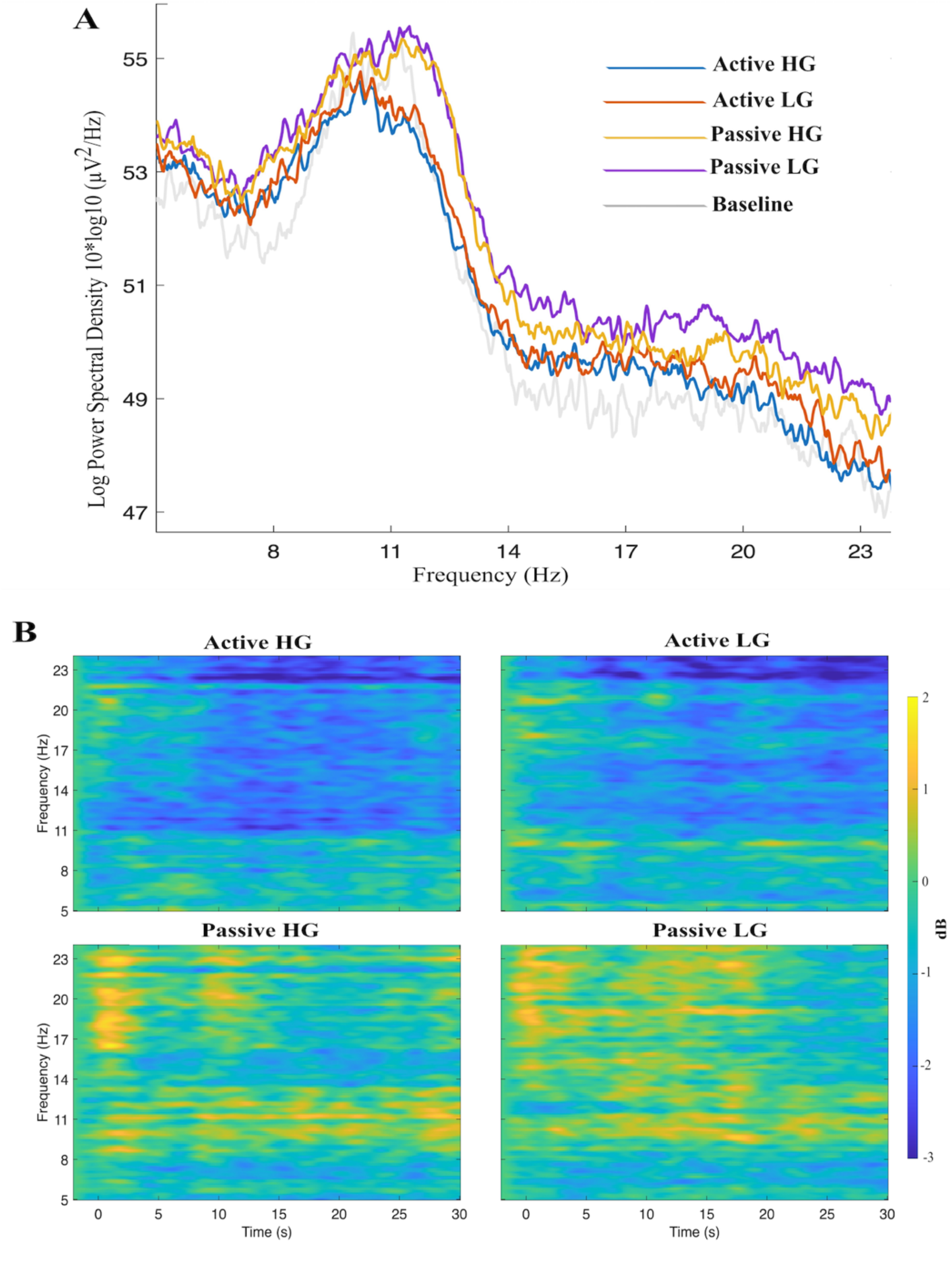
Spectral and ERSPs analyses of electrode C4. **A:** Mean log power spectra: power amplitude (reported in Log Power -y axis) at specific frequency bands (alpha and beta -x axis) for the four experimental conditions + baseline. µ-related activity in the alpha band was detected at ∼ 11-12 Hz; beta-band peaks (∼19 Hz) comprise µ harmonic and possibly other beta-band activity. **B**: Time-frequency plots: ERD (in blue) and ERS (in yellow) for alpha and beta (y axis) during the epoch duration (-2 to 30 sec from the stimulus onset -x axis).

**Appendix Fig. 4.**
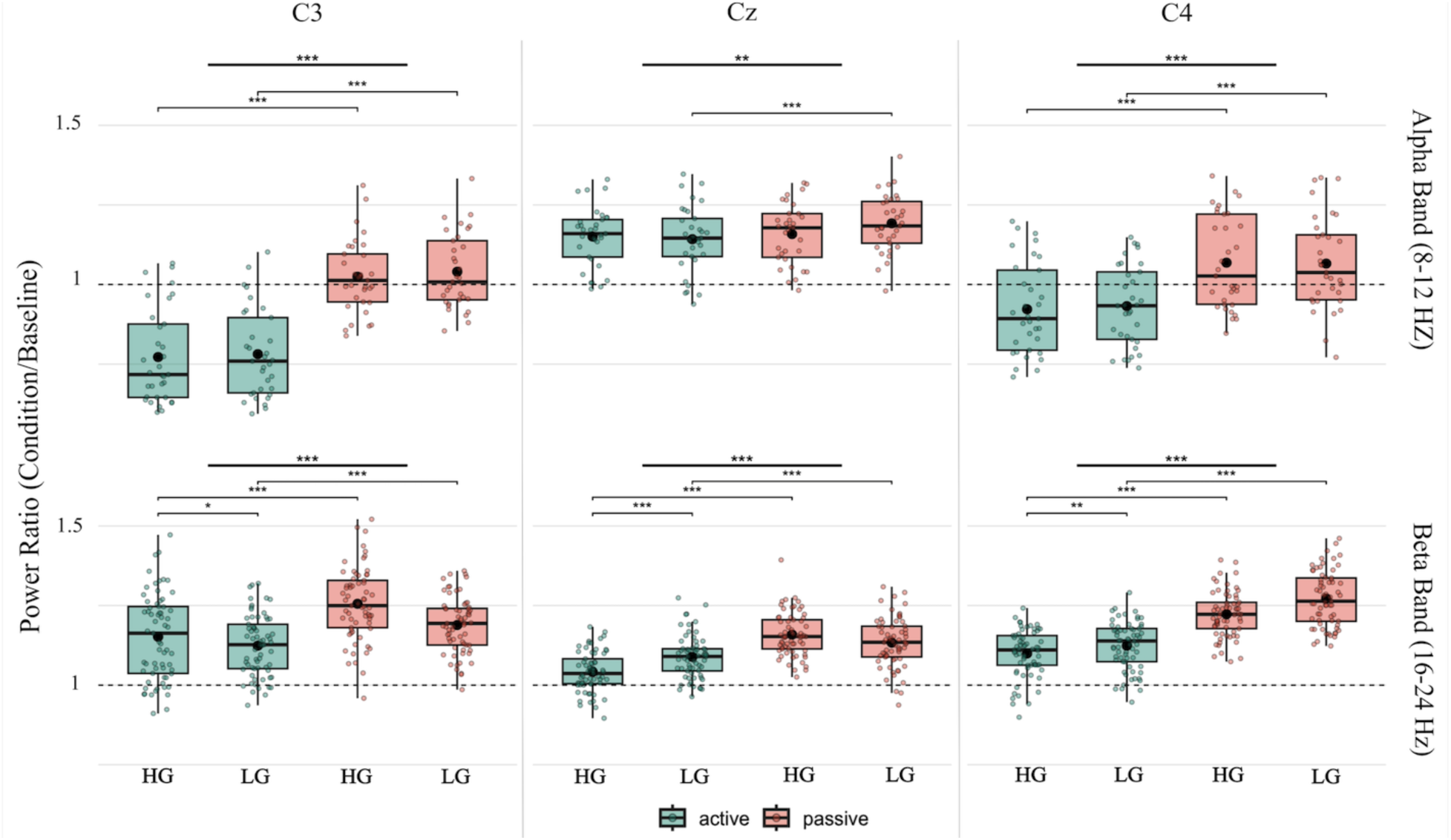
Ratios of mean power during active HG, active LG, passive HG and passive LG conditions to mean power in the baseline (ERD, this ratio < 1; ERS, this ratio > 1) for channels (columns) at specific frequency bands (alpha - first row; beta - second row). Ticker lines at the top mark significant differences between tasks (active in green vs passive in pink); thin lines mark significant differences between pairs of conditions from the post hoc comparisons.

**Appendix Fig.5.**
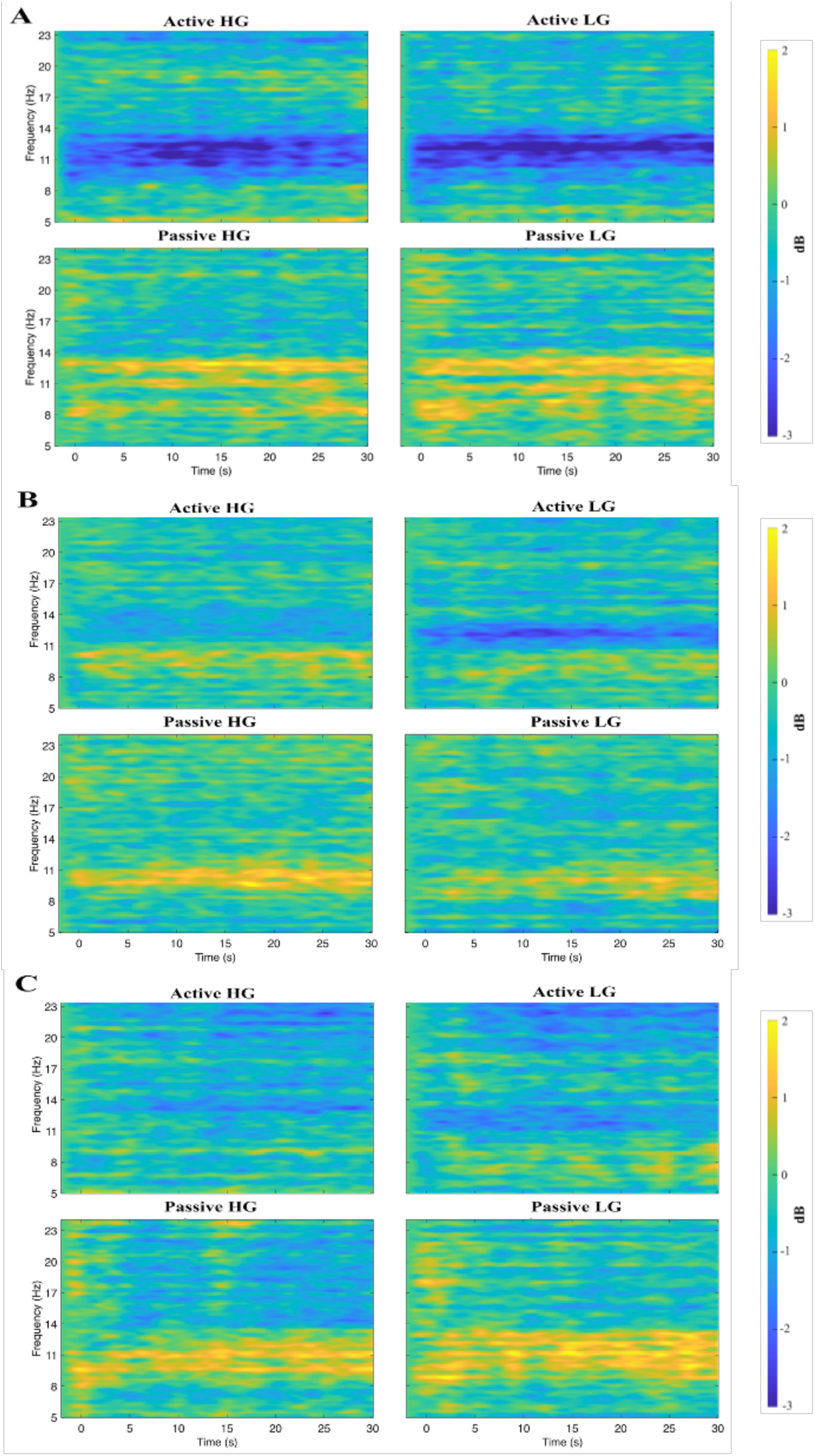
ERSPs analyses of Clusters. Time-frequency plots: ERD (in blue) and ERS (in yellow) for left (**A**) midline (**B**) and right (**C**) clusters for alpha and beta (y axis) during the epoch duration (-2 to 30 sec from the stimulus onset -x axis).

